# Sexual dimorphism in mastoid process volumes measured from 3D models of dry crania from medieval Croatia

**DOI:** 10.1101/2021.01.04.425320

**Authors:** Anja Petaros, Sabrina B. Sholts, Mislav Čavka, Mario Slaus, Sebastian K.T.S. Wärmländer

## Abstract

3D analysis of skeletal volumes has become an important field in digital anthropology studies. The volume of the mastoid process has been proposed to display significant sexual dimorphism, but it has a complex shape and to date no study has quantified the full mastoid volume for sex estimation purposes. In this study we compared three different ways to isolate the volume of the mastoid process from digital 3D models of dry crania, and then evaluated the performance of the three different volume definitions for sex estimation purposes. A total of 170 crania (86 male, 84 females) excavated from five medieval Croatian sites were CT-scanned and used to produce 3D stereolitographic models. The three different isolation techniques were based on various anatomical landmarks and planes, as well as the anatomy of the mastoid process itself. Measurements of the three different mastoid volumes yielded different accuracies and precisions. Interestingly, anatomical structures were sometimes more useful than classical landmarks as demarcators of mastoid volume. For all three volume definitions, male mastoid volumes were significantly larger than female volumes, in both relative and absolute numbers. Sex estimation based on mastoid volume showed a slightly higher precision and better accuracy (71 % correct classifications) than visual scoring techniques (67 %) and linear distance measurements (69 %) of the mastoid process. Sex estimation based on cranial size performed even better (78 %), and multifactorial analysis (skull size + mastoid volume) reached up to 81% accuracy. These results show that measurements of the mastoid volume represent a promising metric to be used in multifactorial approaches for sex estimation of human remains.

## 1. INTRODUCTION

The mastoid process and its surrounding region represent one of the most sexually dimorphic parts of the human skull, and is often included in data collection protocols and sex estimation methods (Buikstra and Ubelaker, 1994; Garvin et al., 2014; Jung and Woo, 2016; Langley et al., 2017; Lewis and Garvin, 2016; Nagaoka et al., 2008; Ramsthaler et al., 2010; Rogers, 2005; Stevenson et al., 2009; Walker, 2008; Williams and Rogers, 2006; Yilmaz et al., 2015). The mastoid process can be assessed either visually by its massiveness and voluminosity (Buikstra and Ubelaker, 1994; Walker, 2008), or metrically by its length (sometimes referred to as height), which typically is measured as the vertical projection of the mastoid process below and perpendicular to the Frankfurt horizontal (FH) plane (Moore-Jansen and RL Jantz, 1986; Buikstra and Ubelaker, 1994; Moore-Jansen, RL Jantz, Ousley SD, 1994, Howells 1973). While visual techniques are quick and economical, metric techniques are currently preferred in bioarcheological and forensic anthropological work, as they are less subjective, better suited for statistical analysis, and typically evaluate skeletal traits with higher accuracy (Garvin and Ruff, 2012; Shearer et al., 2012). The mastoid process might be an exception, since in many occasions visual techniques have performed better than metric ones in terms of accuracy and precision (Petaros et al., 2015). This is likely related to the complex shape of the mastoid process, which may display more sexual dimorphism than its size. The mastoid furthermore lacks well-defined anatomical borders, and there are no convenient landmarks to use that directly reflect mastoid size.

Measuring the distance between *mastoidale* and *porion* becomes rather subjective (Petaros et al., 2015; Saini et al., 2012), as this measurement depends on the orientation/angulation of the mastoid process and the distance between the mastoid tip and the observer (Petaros et al., 2015). The difficulties involved in measuring the mastoid process have been noted by Howells as early as in 1973 (Howells, 1973), and later on also by Nagaoka (2008), Petaros et al (2015), and Langley et al (2016). As a response, the latter group of authors proposed a new and possibly more precise approach for measuring mastoid length (height) (Langley et al., 2016; Langley et al., 2018), which now is included in the manual for standardized recording procedures (Langley et al., 2016). Recently, however, even the precision and accuracy of visual techniques for mastoid evaluation have been questioned: they may perform worse than previously thought, and may significantly depend on the population studied (Lewis and Garvin, 2016).

Despite the limitations mentioned above, the mastoid process often is used in anthropological and bioarchaeological studies that evaluate cranial sexual dimorphism, either by validating existing methods on different populations (Buran et al., 2018; Franklin et al., 2005b; Galdames et al., 2008; Gangrade et al., 2013; Jaja et al., 2013; Kanchan et al., 2013; Kittoe et al., 2012; Madadin et al., 2015; Manoonpol and Plakornkul, 2012; Sujarittham et al., 2011) or developing new ones (Abdel Fatah et al., 2014; Amin et al., 2015; de Paiva and Segre, 2003; Jung and Woo, 2016; Langley et al., 2017; Nagaoka et al., 2008; Sharma et al., 2013; Stevenson et al., 2009; Sumati et al., 2010). Mastoid length is also among the features included in the FORDISC software for sex/ancestry group classification (Jantz and Ousley, 2005). The large amount of studies using the mastoid process have brought to light another problem that makes comparisons between different studies difficult, namely that of a large variation in the landmarks chosen for taking the measurements, as well as a significant inconsistency in the terminology used to describe the measurements, (Petaros et al., 2015). This may explain why researchers sometimes report lower precisions for mastoid metrics, and often fail to validate previously proposed methods for mastoid measurements (Jaja et al., 2013; Kemkes and Gobel, 2006; Nagaoka et al., 2008). In Tables 1 and 2, we list the results of a large number of earlier studies that have investigated the sexual dimorphism of the mastoid process in different populations and samples using the same approaches.

**Table 1.**
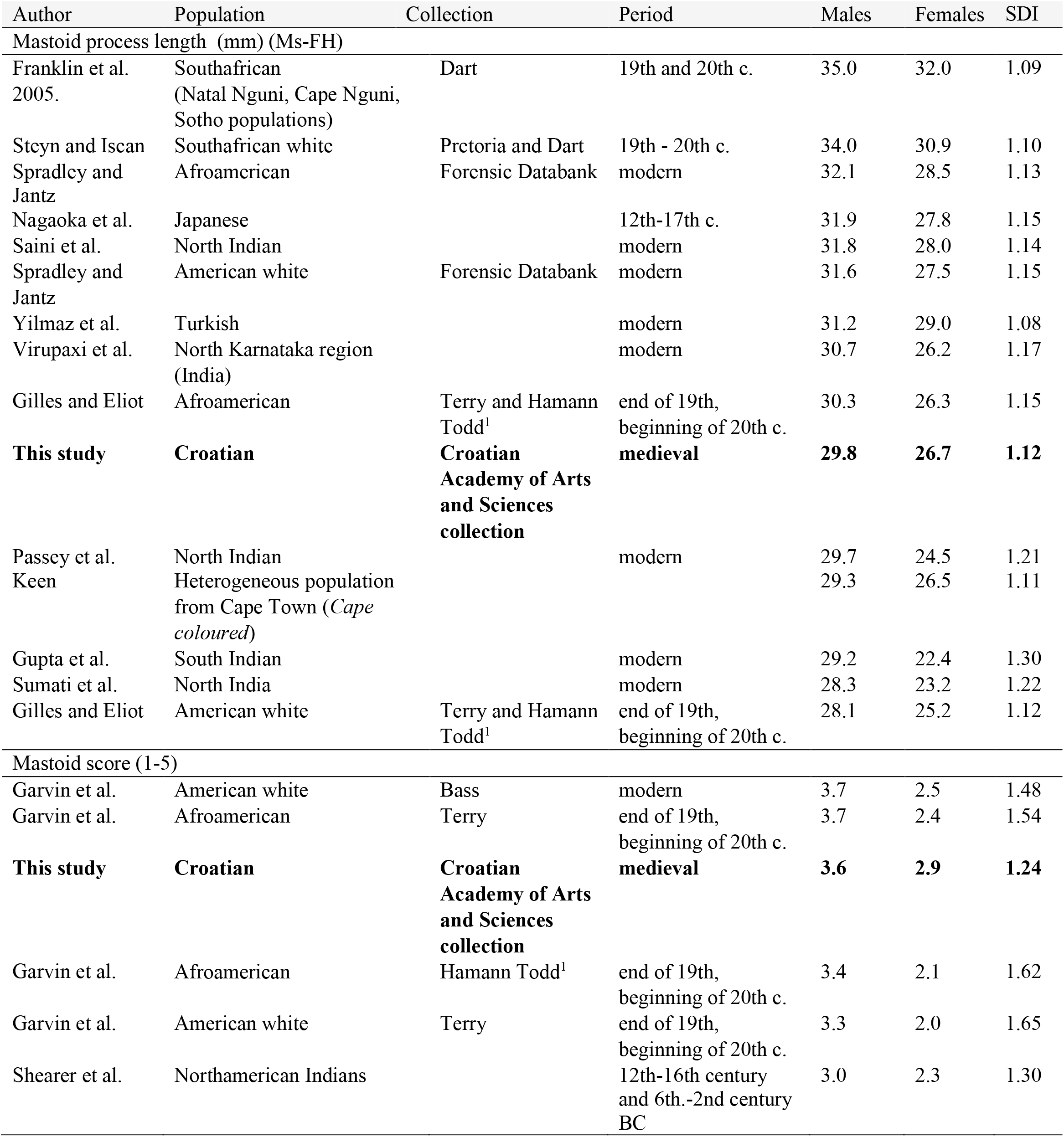
Comparison of traditional mastoid measurements recorded for different world populations from different time periods. Ms= mastoidale, FH= Frankfurt horizontal. ^1^Hamann Todd is composed mainly of the 2nd generation of “Central European” immigrants.

**Table 2.**
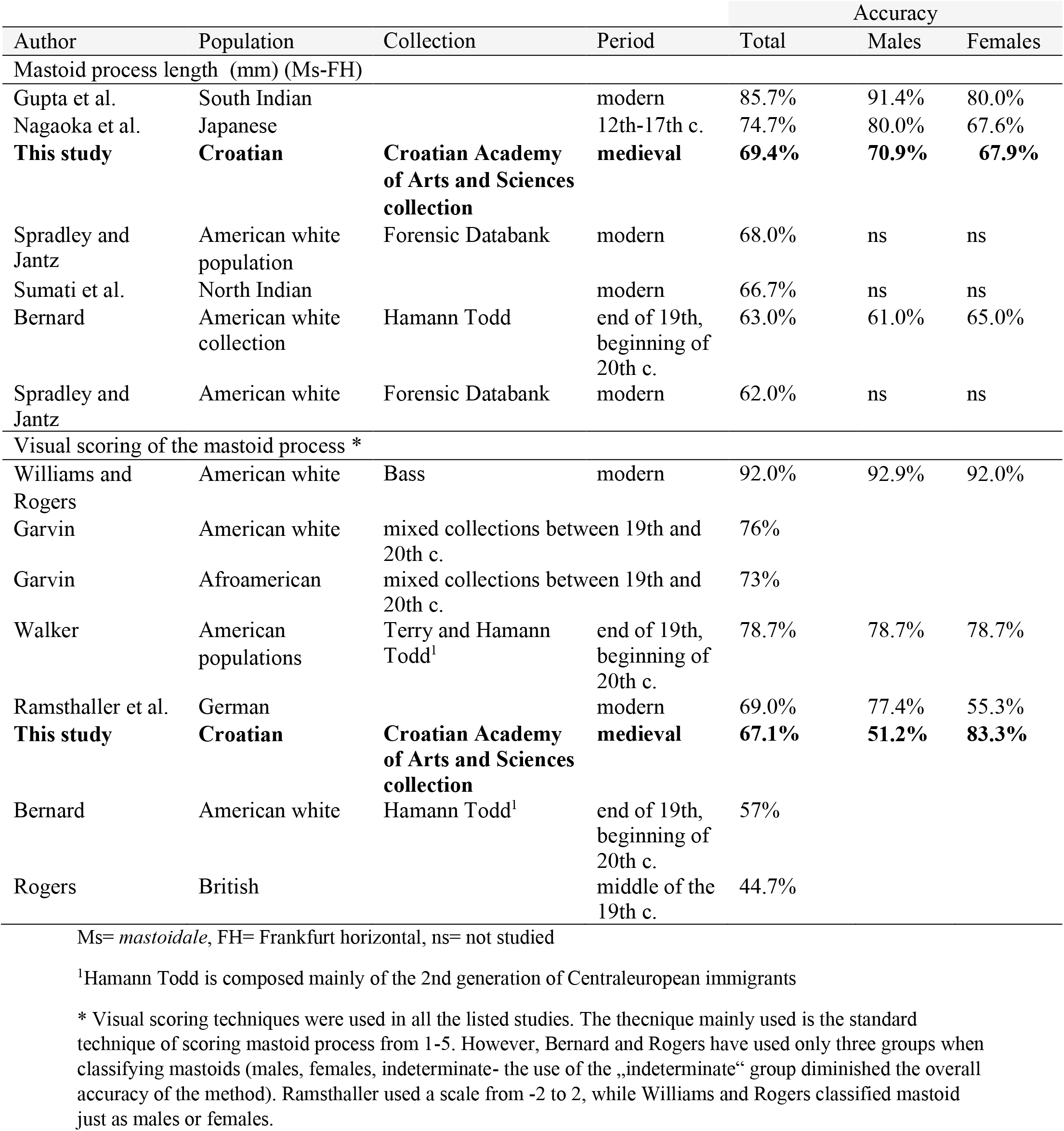
Accuracy differences in using traditional mastoid measurement observed between different populations and collections

Because of the problems involved in measuring the mastoid process, a number of novel approaches have been presented. Although some of them facilitate the quantification of morphological features, only a few address the issue of mastoid shape. Jung and Woo (2016) used geometric morphometrics analysis of sliding landmarks to investigate mastoid shape and centroid size, while Abdel Fatah et al. (2014) used an innovative 3D approach that by capturing primary shape variation in the skull, demonstrated sexually dimorphic shape differences in the mastoid process. Other studies have investigated classical linear distances and angles (Buran et al., 2018; Kramer et al., 2018; Yilmaz et al., 2015; Zaafrane et al., 2018), or the area of the “mastoid triangle”: porion - asterion – mastoidale) (de Paiva and Segre, 2003; Gangrade et al., 2013; Ibrahim et al., 2018; Jain et al., 2013; Madadin et al., 2015; Toneva et al., 2019). Baki Allam and Baki Allam (2016) presented a novel approach to mastoid shape and volume analysis by demonstrating sexual dimorphism in a mastoid volume defined by (mastoid height × maximal oblique sagittal diameter × maximal oblique coronal diameter × 0.52). To the best of our knowledge, however, no one has so far analyzed the mastoid from its full 3D volume (Petaros et al., 2015).

In this study we test three different approaches for the quantification of the mastoid volume based on the osteometrical points in the nearby regions as well as the anatomy and embryology of the mastoid region. Further, we investigate if the objectively defined 3D volumes encompassing different parts of the mastoid process can be used for statistical sex estimation, and discuss the problem of how to best delineate such a volume.

## 2. MATERIAL AND METHODS

### The sample and related limitations

The sample for this study consists of 170 medieval crania (86 males and 84 females) from the osteological collection of the Croatian Academy of Arts and Sciences (Zagreb, Croatia). The crania originate from five medieval archaeological sites in the Dalmatian region of southern Croatia: Velim, Radašinovci, Dugopolje, Koprivno, and Šibenik (Figure 1; Table 3). The first two date to the Croatian Early Medieval Period, i.e. 9^th^ – 11^th^ c., while the last three to the Late Medieval Period, i.e. 11^th^-16^th^ c. (Premuzic, 2013; Slaus et al., 2011). All crania belong to well-preserved and complete adult skeletons, where adulthood was determined from the fusion of long bones and eruption of a third molar (Slaus, 1997; Slaus et al., 2003). Sex was determined from visual inspection of the pelvis and anthropometry of long bones.

**Figure 1.**
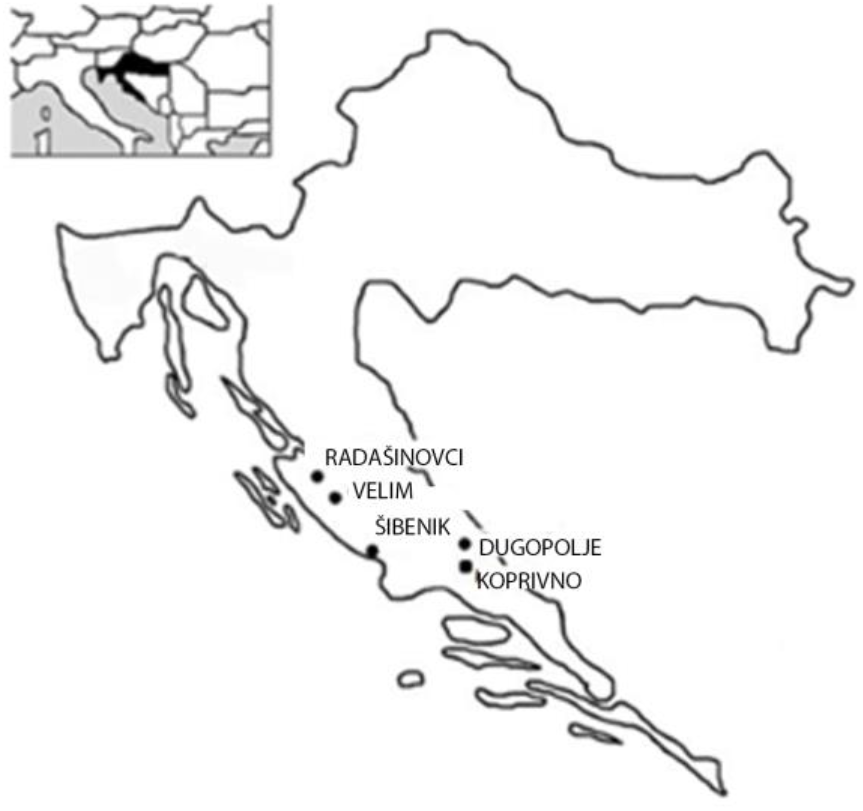
Map of Croatia showing the Medieval archaeological sites where the skeletons used in the study were excavated.

**Table 3.**
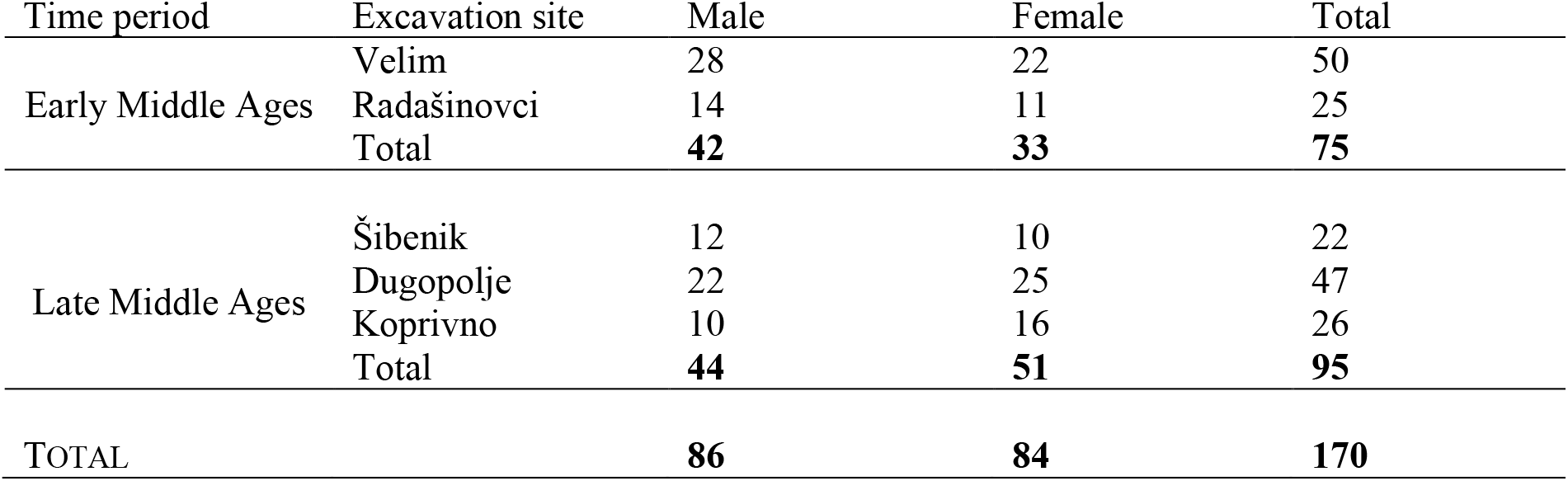
Distribution of the sample with regard to sex, excavation site, and time period

All crania selected for the study were without antemortem head trauma or significant damage to the mastoid regions, and they all had well-defined Frankfurt horizontal planes, i.e. the *orbitale* and both *porion* landmarks were always present. The study primarily evaluated left mastoid processes (n = 151), but right mastoid processes were analyzed when the left one was missing or damaged (n = 19).

A sample of well-preserved archaeological crania was chosen for the study, as it allows recording of 3D models with higher resolution than e.g. CT scans of living patients. At the time of the study, this was the only sample in the territory available for an extensive radiologic and anthropological analysis. The studied crania belong to a reference collection previously used to develop sexing standards for the Croatian medieval population (Slaus, 1997). The skeletons in this collection are unidentified, and thus sex is not known but estimated from a range of skeletal features such as long bone size. Although the anthropological sexing accuracy of complete skeletons go well above 90% (de Paiva and Segre, 2003; Sjøvold, 1988), and although the anthropological sexing of skeletons from medieval archaeological sites in the Dalmatian region has been verified by DNA analysis (Basic et al., 2013), we cannot exclude the possibility that some skulls may have been wrongly categorized. This is a limitation that should be taken in consideration when evaluating the presented results.

### Traditional visual and metrical analysis of the mastoid process

Each mastoid process was evaluated both visually and metrically following the protocols in the *Standards for Data Collection from Human Skeletal Remains*” (from here on “*Standards*”) (Buikstra and Ubelaker, 1994). Visual analysis was conducted by placing each cranium on its right side and assigning to the relative size/volume of the mastoid process a number ranging from 1 (minimal expression) to 5 (maximal expression). Measurements of the mastoid process length (from *porion* to *mastoidale*) were performed on digital 3D models to eliminate errors originating from orientation and mastoid-observer distance (Casado, 2017) (Fig 2F).

**Figure 2.**
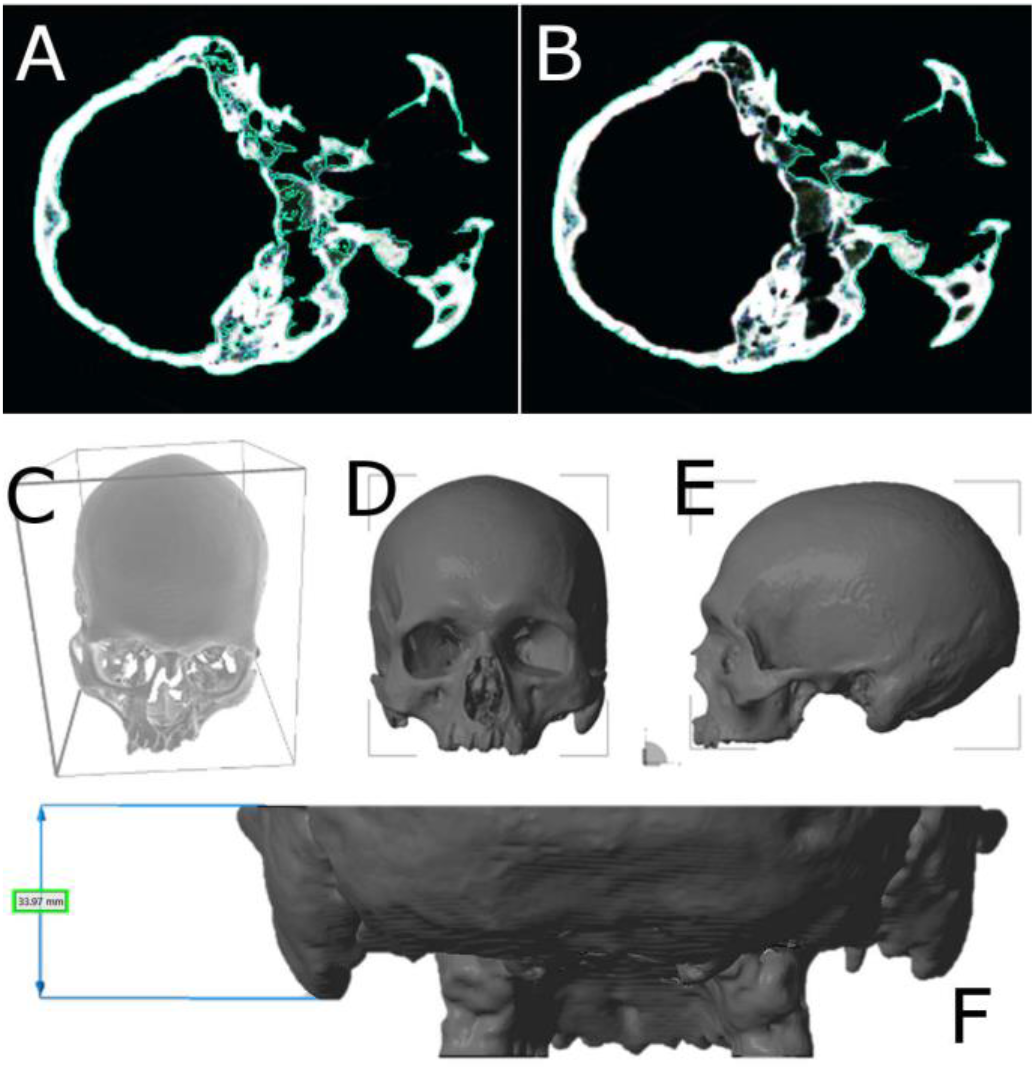
A-B) Two different ways of segmenting the skull, using 3D CT models and two different functions in the 3D doctor software. A) segmentation of the skull external outline only. B) segmentation of all boundaries. In the study, segmentation was performed just using the outline contour of the skull. C-E) Illustrations of the bounding box used to calculate the volume of each skull. See Shearer et al. (2012) for a full discussion of this concept. F) Measurement of the mastoid length performed on digital 3D models using the Netfabb software

### 3D imaging

#### Creating 3D models of the crania

The crania were scanned at the Department of Diagnostic and Interventional Radiology, University Hospital Dubrava, Zagreb, Croatia, with an MDCT unit (Sensation 16, Siemens AG Medical Solutions, Erlangen, Germany) operating at 120 kV/320 mA and recording continuous layers without overlap, using 12 x 0.75 mm collimation. The resulting DICOM data files (approximately 250 slices per cranium) were imported into the 3D Doctor imaging program (Able Software Corp., 1998-2011). A soft tissue kernel was used for CT image reconstruction, followed by threshold-based semi-automatic bone extraction. The segmentation was done using only the external outline of the skull (boundary type: outline only function; Figures 2A-B), thus circumventing the complex inner architecture of the temporal bone. In this way, the volume of the mastoid air cells and the thickness of the cranial bones (which are under effect of environmental factors and sex, as well as subject to idiosyncrasy) were excluded from the analysis, and the mastoid process was considered to be a solid object – similar to how it is perceived during anthroposcopical inspections.

After completing the segmentation, all cranial 3D models were reconstructed using the surface-rendering technique, keeping the maximum denseness of the triangle mesh, and exported as stereolitography (STL) models retaining maximum model resolution. The Netfabb Studio Professional software (netfabb GmbH, Germany, 2009) software was then used to orient and level (using x,y,z coordinates) the 3D models along the Frankfurt Horizontal plane. The leveling was done to avoid any uneven points between the right and left side of the skull. Although the measured volumes of cranial 3D models to some extent depend on the resolution and parameters used to create and process the 3D models (Sholts et al., 2010), using identical settings throughout the study ensures that comparisons are not biased by the 3D model processing.

#### Volume measurements from cranial 3D models

The Netfabb software was employed to digitally isolate the mastoid process and its surrounding region(s) from the cranial 3D models. Here, we consider the mastoid process to consist of two different parts separated by the petrosquamous suture: the antero-superior squamous part and the postero-inferior petrous part (Mansour et al., 2013). We used three different approaches to isolate the mastoid process, employing different geometric planes defined *via* anatomical and anthropological landmarks and structures (Figure 3).

**Figure 3.**
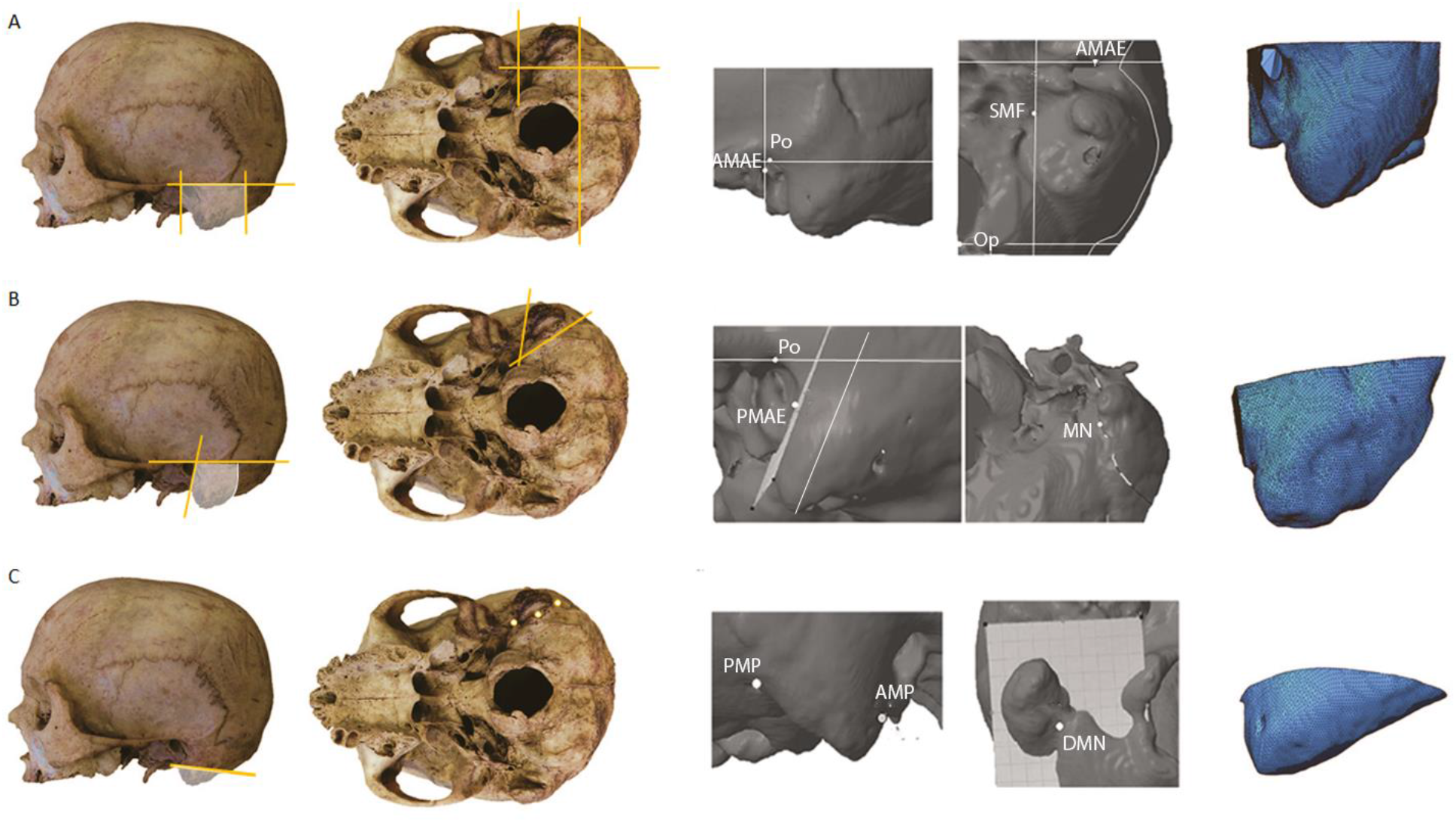
Planes and landmarks used to isolate the three different mastoid volumes presented in the skull profile and inferior view: A. Isolation of the wider mastoid region (volume 1) – AMAE-anterior margin of the *meatus acusticus externus*, Op- opisthion, SMF- stylomastoid foramen; B. Isolation of the mastoid in the narrower sense (volume 2) – PMAE- posterior margin of the *meatus acusticus externus*, Po-porion, MN- mastoid notch - anterior mastoid process plane; C.) Isolation of the mastoid tip (volume 3) – AMP – anterior root of the mastoid process, DMN- deepest point of the mastoid notch, PMN-posterior end of the mastoid notch.

Volume 1 (“Vol-1”) comprises the broader mastoid region, i.e. the mastoid process and parts of the temporal and occipital bones. This volume is defined by four cutting planes: one transverse plane passing through *porion* which cuts the region superiorly (Frankfurt horizontal plane), and three sagittal planes passing through respectively the anterior border of the external auditory meatus-cutting the region anteriorly, the stylomastoid foramen – cutting the region medially, and *opisthion –* cutting the region posteriorly (Figure 3A). The landmark *asterion* was avoided when defining this volume, as its location is idiosyncratic and also depends on age and population affinity (Day and Tschabitscher, 1998; Mwachaka et al., 2010; Ross et al., 1999; Sripairojkul and Adultrakoon, 2000), the position of the opisthion itself can vary between populations being part of the cranial base and dependent on the shape and size of the foramen magnum. Moreover since foramen magnum size differs between sexes, Volume 1 reflects not only the “proper” mastoid process but also some neighbouring regions and anatomical parts. The idea behind Vol-1 was to investigate a volume that is defined by traditional well-defined anthropometric points, and which encompasses the mastoid process in its entirety.

Volume 2 (“Vol-2”) comprises the mastoid process in the narrow sense, i.e. the squamous and petrous portions of the mastoid process without any other part of the temporal bone. Vol-2 is delineated by three planes (Figure 3B): transverse plane passing through *porion* (FH plane), a plane parallel to the directional axis of the mastoid and passing through posterior border of the external auditory meatus, and an oblique plane passing through the mastoid notch (i.e., the digastricus muscle attachment site). In difference from Vol-1, Vol-2 is defined mainly by anatomical structures related to the mastoid region that are not considered traditional anthropometric landmarks.

Volume 3 (“Vol-3”) comprises the mastoid tip, i.e. the bulging part of the mastoid process that protrudes underneath the temporal bone. This volume was isolated using a single plane defined by three non-traditional landmark points located on the mastoid process itself (Figure 3C): the point where the mastoid process detaches from the temporal bone (anterior point), the posterior end of the mastoid notch (posterior point), and the deepest point of the mastoid notch (medial point). This volume does not consider the external auditory meatus (EAM), in line with the suggestions for the visual inspection standards (Buikstra and Ubelaker, 1994). After cutting along the delineation planes, the remaining 3D model part sometimes contained additional bone not connected to the mastoid, which was manually removed using the Netfabb software

For each 3D model, all three volumes were calculated using a function integrated in the Netfabb software.

### Relative volume calculation

As adult males generally have larger skulls than adult females, relative mastoid volumes were calculated following the protocol developed by Shearer and al. (Shearer et al., 2012), i.e. by dividing the mastoid volume with the corresponding cranium’s 3D bounding box volume (defined as the product of the height, width, and depth of the box surrounding the 3D model; these measurements are routinely obtained from 3D software). Because some skulls had lost teeth postmortem or were edentulous, the lower border of the bounding box was defined by the lower border of the maxilla rather than the teeth (Figures 2C-E).

### Intra-observer error

To test the intra-observer precision for each of the three volumes, the same observer (AP) re-evaluated 50 randomly selected crania. Except for the CT scanning, all stages of the 3D model analysis described above were repeated a second time, i.e. segmentation, construction, and orientation of the STL models, as well as isolation of the three different mastoid volumes.

### Statistical analysis

Intra-observer precision for each of the three volumes was evaluated by calculating coefficients of variability. Sexual dimorphism was evaluated using student’s t-test and comparisons of the sexual dimorphism index (“SDI”), defined as the mean value for males divided by the mean value for females. The accuracy of each method for sex estimation was tested using discriminant function analysis. Multifactorial analysis was performed to investigate whether involving more traits would produce a higher classification accuracy. The relation between mastoid volume and mastoid length or mastoid score was evaluated via correlation coefficients (Pearson’s r for the first relation; Spearman’s rank correlation coefficient (rho) for the latter). For all analyses, results with p < 0.05 were considered statistically significant.

## 3. RESULTS

### Intra-observer precision of the cutting methods

Differences between the first and the second measurement for the 50 randomly selected crania range between 0.03 to 1.38 cm^3^ (median 0.3 cm^3^) for Vol-1, from 0.01 to 0.64 cm^3^ (median 0.18 cm^3^) for Vol-2, and from 0.0 to 1.01 cm^3^ (median 0.09 cm^3^) for Vol-3. The mean, minimum, and maximum values of the coefficient of variability (CV) are shown in Table 4.Volumes 1 and 2 and display the highest precision, with mean CV values below 2.5%. Volume 3 shows poor repeatability, with a mean CV value around 8% and occasional CV values reaching 51%.

**Table 4.**
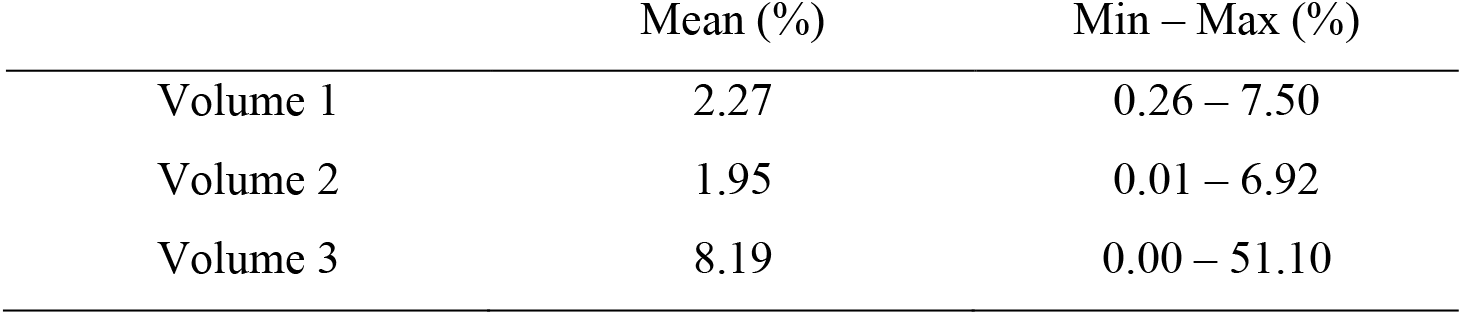
Mean, minimum, and maximum differences for the coefficient of variation for the three studied mastoid volumes

### Sexual dimorphism in the mastoid 3D volumes

For all three absolute volumes, male values are on average significantly larger than female (Figure 4 and Table 5). There is also a tendency for male mastoid volumes to be larger in the Late Middle Ages than in the Early Middle Ages, suggesting possible secular changes (Figure 4).

**Figure 4.**
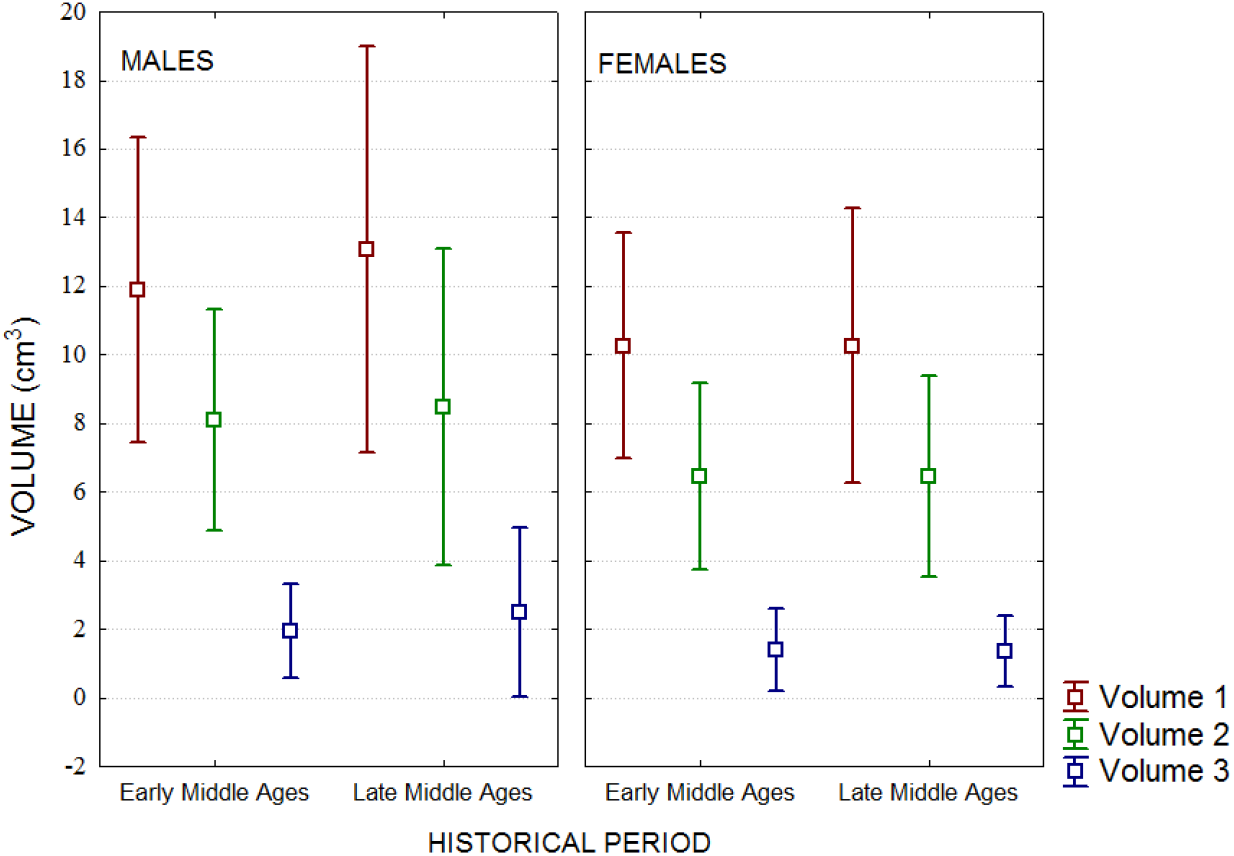
Volume differences between the sexes and between different historical periods.

**Table 5.** Male and female volume, lenght and score values with corresponding male-female differences, statistical significance and sexual dimorphism index

|  | Male |  | Female |  | Difference<br>(cm<sup>3</sup>) | T<br>value | P value | SDI |
| --- | --- | --- | --- | --- | --- | --- | --- | --- |
|  | Mean<br>(cm<sup>3</sup>) | SD<br>(cm<sup>3</sup>) | Mean<br>(cm<sup>3</sup>) | SD<br>(cm<sup>3</sup>) |  |  |  |  |
| Volume 1 | 12.50 | 2.67 | 10.26 | 1.85 | Δ2.24 | 6.31 | <0.0001* | 1.22 |
| Volume 2 | 8.29 | 1.99 | 6.45 | 1.41 | Δ1.84 | 6.97 | <0.0001* | 1.28 |
| Volume 3 | 2.23 | 1.04 | 1.37 | 0.55 | Δ0.86 | 6.63 | <0.0001* | 1.63 |
|  | Mean<br>(‰**) | Range<br>(‰) | Mean<br>(‰) | Range<br>(‰) |  | T<br>value | P value | SDI |
| REL V1 | 0.28 | 0.16-0.45 | 0.25 | 0.15-0.34 |  | 2.73 | 0.007* | 1.12 |
| REL V2 | 0.18 | 0.09-0.34 | 0.16 | 0.09-0.29 |  | 3.72 | 0.0003* | 1.13 |
| REL V3 | 0.05 | 0.01-0.12 | 0.03 | 0.003-0.06 |  | 5.17 | <0.0001* | 1.67 |
|  | Mean | SD | Mean | SD |  | T (Z)<br>value | P value | SDI |
| Length | 29.81 mm | 3.02 mm | 26.65 mm | 2.87 mm |  | 6.99 | <0.0001* | 1.12 |
| Score | 3.60 | 0.82 | 2.87 | 0.77 |  | 5.02 | <0.0001* | 1.25 |

\* statistically significant
\*\* the results of the division of the mastoid volume with the corresponding cranium's 3D bounding box volume are presented in per mille, according to Shearer et al (2012).
SD= standard deviation, SDI= sexual dimorphism index

Sex estimation accuracies achieved by absolute volumes are included in table 6, and range between 66,5 and 71,2%. For all three volumes females are classified with higher accuracy than males, which is to be expected as female mastoid volumes display less variation than male ones (Table 5).

**Table 6.** Sex estimation based on different parameters ^a^ Bounding box measurement is described in the materials and methods section

|  | Total (n=170) (%) | Male (n=86) (%) | Female (n=84) (%) |
| --- | --- | --- | --- |
| VOLUMES |  |  |  |
| Absolute volume values |  |  |  |
| Volume 1 | 66.5 | 65.1 | 67.9 |
| Volume 2 | 70.6 | 68.6 | 72.6 |
| Volume 3 | 71.2 | 67.4 | 75.0 |
| Relative volume values |  |  |  |
| Volume 1 | 60.2 | 61.4 | 59.0 |
| Volume 2 | 62.1 | 62.6 | 61.5 |
| Volume 3 | 68.9 | 67.5 | 70.5 |
| TRADITIONAL MEASUREMENTS |  |  |  |
| Mastoid length | 69.4 | 70.9 | 67.8 |
| Mastoid score | 67.0 | 51.2 | 83.3 |
| SKULL SIZE |  |  |  |
| Bounding box<sup>a</sup> | 78.4 | 73.5 | 83.3 |
| COMBINATION |  |  |  |
| Volume 1+ length | 69.4 | 66.3 | 72.6 |
| Volume 2+ length | 69.4 | 67.4 | 71.4 |
| Volume 3+ length | 74.1 | 74.4 | 73.8 |
| Volume 1+ score | 68.2 | 64.0 | 72.6 |
| Volume 2+ score | 72.6 | 67.6 | 62.8 |
| Volume 3+ score | 74.1 | 68.6 | 79.8 |

When relative rather than absolute volumes are evaluated, in order to control for sexual dimorphism in overall skull size, the male/female difference is still evident but less clear: lower SDI values and higher p values are obtained (Table 5). The accuracy results (Table 6) show that relative volumes perform around 7% worse than estimations based on absolute volumes.

### Traditional mastoid measurements: length and scoring

The traditional anthropological techniques of length measurement and visual scoring of the mastoid process reveal sexual dimorphism at highly significant levels, i.e. p < 0.0001 for both parameters (Table 5). The average sexing accuracies are 69,4% for mastoid length and 67% for mastoid scores (Table 6). While the accuracies based on length are rather similar for males (70.9%) and females (67.8%), the accuracies based on visual scoring display a distinct male/female discrepancy (i.e. 51.2%/83.3%).

The traditional measurements showed a significant correlation with all three mastoid volumes, but not with overall skull size (Table 7). The strongest correlations were obtained for mastoid length with absolute volumes. When mastoid volume is combined with mastoid score to produce discriminant functions for sex estimation, slightly better results are achieved compared to using either parameter alone (i.e. up a few percent; Table 6). Small improvements in sex estimation accuracy are obtained also when mastoid volume is combined with mastoid length, although the combination of length and Vol-2 yielded no improvement at all (Table 6).

**Table 7.** Correlations between traditional mastoid measurements and mastoid absolute and relative volumes. For the length and skull size, Pearson coefficient r values are shown, while Spearman coefficient R values are shown for the mastoid scores.

|  | Absolute volume |  | Skull size<sup>a</sup> | Relative volume |  |
| --- | --- | --- | --- | --- | --- |
|  | Length | Mastoid scores |  | Length | Mastoid scores |
| Volume 1 | 0.73* | 0.53* | 0.31 | 0.58* | 0.40* |
| Volume 2 | 0.76* | 0.61* | 0.30 | 0.64* | 0.51* |
| Volume 3 | 0.58* | 0.51* | 0.25 | 0.53* | 0.46* |

\* statistically significant
<sup>a</sup>The skull size was measured as the cranium bounding box, as described in the materials and methods section

## 4. DISCUSSION

### Precision of the three different volumes

The mastoid process represents a complex region in the human skull that lacks clear landmarks points to delineate it. It is therefore a challenging part of the human skull to precisely isolate for 3D studies. In this research, we compared three different ways to isolate the volume of the mastoid process from digital 3D models.

Vol-2 displays the highest precision, with a CV < 2% (Table 4). Similar high precisions have previously been reported for volume measurements of the mental eminence and glabella (Garvin and Ruff, 2012). Because a main source of error in digital anthropology is locating landmarks (Shearer et al., 2012), the good repeatability of Vol-2 is likely related to its three cutting planes being defined not only by traditional landmarks, but also by anatomical locations such as the anterior mastoid edge and mastoid notch (Figure 3).

Vol-1 is defined mostly by EAM landmarks, the location of which have shown some population and age dependence (Schulter, 1976). In particular, the achieved a slightly worse (CV around 2.3%, Table 4), which may be related to the difficulties in locating the stylomastoid foramen in the CT models. Due to the segmentation approach that was used (i.e. outline only) the contours of the foramen were not distinct in every 3D model (Figures 2A-B). Earlier work has already noted that certain landmarks that are clearly visible on dry skulls may be difficult to locate on 3D models (Sholts et al., 2011b). Replacing the stylomastoid foramen with another landmark, such as the lateral border of the foramen magnum, might increase the precision of Vol-1.

The worst precision is found for Vol-3, which has a CV around 8.2% (Table 4). This is likely related to the manual cutting of the extraneous bone (in the digital 3D model) and the use of landmarks that are difficult to locate in a reproducible manner. It has previously been reported that the landmarks PEIM and the anterior border of the mastoid process can be located with good precision (Gonzalez et al., 2011). This suggests that the landmark responsible for the large variability of Vol-3 is the deepest point of the mastoid notch, which would be consistent with previous reports of poor precision in landmarks related to the mastoid notch (Howells, 1973; Nagaoka et al., 2008).

Overall, this indicates that volumes 1 and 2 are acceptable for use in anthropological practice, but not Vol-3. Because volumes 1 and 2 require the skull to be in the FH plane, they are both limited by the completeness of the skull, as these volumes only can be calculated for skulls where the landmarks *orbitale* and *porion* are present.

It has often been mentioned that geometric features used in morphometric analysis – such as distances, areas, and volumes – should be defined via landmarks that are classified as Type 1 in Bookstein’s typology. The argument is that the locations of such landmarks can be identified with higher accuracy and precision than the locations of Type 2 and Type 3 landmarks. It has however previously been shown that the landmark measurement precision depends on the method used, e.g. if landmarks are measured from dry bones with calipers or a microscribe, or if digital tools are used to locate landmarks on 3D models (Sholts et al., 2011a). Thus, it has been argued that the preference for Type 1 landmarks in morphometric studies should be re-evaluated, and that Bookstein’s landmark typology may be less useful for research design, especially when digital techniques are involved (Wärmländer et al., 2019). In line with this, we do not here consider which Bookstein Type the employed landmarks might belong to. Instead, we focus on the specific properties of the landmarks in question, and ask if they are suitable for the particular task at hand (i.e., if they are useful for defining mastoid 3D volumes).

### Sexual dimorphism

All three mastoid volumes showed a significant sexual dimorphism in the tested archeological population (p<0.0001; Table 5). The mastoid process follows the growth and development of the human craniofacial complex, which is known to be sex-dependent. Males have a longer growth period that begins later and involves significant and long-lasting growth spurts (Rogers, 1991). Contrary to fast-growing basicranial regions such as the condyles and the foramen magnum, the mastoid process displays a long-lasting growth and consequently reaches a larger size in males than in females (Humphrey, 1998). The mastoid grows in all three dimensions - downwards, backwards, and outwards – and its growth depends in part on the development of air cells inside the mastoid (i.e., pneumatization) (Schillinger, 1939). The latter process likely contributes little to sexual dimorphism, as in adulthood the male/female difference in airspace volume appears to be insignificant (Cinamon, 2009; Hill and Richtsmeier, 2008; Karakas and Kavakli, 2005; Kim et al., 2010). Problems with the air cell development, related to various extrinsic or intrinsic factors such as inflammation, may however interfere with normal mastoid growth. For mastoid sexual dimorphism, muscle activity and biomechanics are likely more important factors: because males have more muscle mass than women and also put more load on their muscles, the stronger tensile forces tend to produce a longer mastoid process in males. Muscle actions are responsible also for the anteromedial inclination of the mastoid process, which is more pronounced in males than in females. Thus, the male/female differences may be primarily expressed through the mastoid tip and the part that serves as attachment site for the *sternocleidomastoideus, longissimus capitis* and *splenius capitis* muscles, while the squamous part might contribute less to the differences between the sexes. Yet, it has been argued that the growth of the mastoid process is largely completed by the age of six (Schillinger, 1939), and it would be interesting to test to what extent muscle action during puberty or in adulthood affects the mastoid shape and size.

Vol-1 is the largest of the studied volumes (Figure 3), and is consequently influenced by all the sex-dependent factors listed above. Yet, its sexual dimorphism index is only a moderate 1.22 (Table 5). Vol-2 only includes the petrous and squamous portions of the mastoid process, and is consequently smaller than Vol-1. While the petrous part serves as an attachment site for neck muscles and thus expresses sexual dimorphism mostly through the length of the mastoid process, the sexual dimorphism of the central squamous part seems to be influenced more by the growth of the temporal bone. This might contribute to a different lateral protrusion and bulging of the mastoid process between males and females. The SDI value for Vol-2 is 1.28, i.e. slightly larger than for volume 1, showing that the larger region captured by volume 1 does not make this volume more sexually dimorphic overall, even though the additional regions themselves are dimorphic. Vol-3 consists solely of the tip of the mastoid process where the digastric muscle attaches (Figure 3). It has a large SDI of 1.63 (Table 5), which in part can be attributed to the sex differences in the locations of the landmarks that define the cutting plane. When PEIM is located more posteriorly and superiorly, the cutting plane will isolate a larger volume of the mastoid tip. The localization of PEIM depends on the shape of the mastoid notch, which is influenced by the activity of the digastric muscle that originates there. A deeper and longer mastoid notch yields a more superior and posterior localization of PEIM, thus contributing to a larger Vol-3. The large SDI of Vol-3 should however be taken with some caution due to the poor repeatability of this volume (see discussion above).

### Sex estimation accuracy

Sex estimation results based on the three mastoid volumes were found to vary between 66.5 and 71.2% (Table 6). Although Vol-3 showed the highest discriminatory power, its practical value is questionable due to its low precision. Vol-1 showed good accuracy and precision, but this volume encompasses not only the mastoid process but also some parts of surrounding regions including the foramen magnum, which itself shows significant sexual dimorphism (Gapert et al., 2009; Madadin et al., 2017; Singh et al., 2017; Uysal et al., 2005). Vol-2 might be the best choice for practical use, given its high precision (1.95 %; Table 4) and good accuracy (70.6 %, Table 6). Vol-2 also better isolates the mastoid process from the rest of the skull, and thus may reflect exclusively the dimorphism of the process itself. The discriminate function analysis better classified females than males (Table 6), which is in line with the lower variation observed for female mastoid volumes (Table 5). In contrast, except when using the highly variable Vol-3, discriminant functions based on relative volume values slightly better classify males (Table 6). This suggests that the larger variation in mastoid size for males is related to a larger variation in overall skull size.

The accuracy analysis furthermore demonstrated that relative volumes performed worse than absolute volumes. This is not unexpected: it is well known that males have larger skulls than females, and the sex estimation accuracy based on skull size alone is 78.4%(Table 6) These results are in line with the observation that combining shape and size parameters generally improves sex estimation (Kimmerle et al., 2008; Gonzalez et al., 2011). Interestingly, although the correlation between mastoid volume and skull size is low and not statistically significant (Table 7), sex estimation based on combined skull size and mastoid volume data is improved up to 81 % (data not shown in tables).

Slightly worse sex estimation is produced by mastoid length (69.4%, Table 7) and mastoid score (67.1%, Table 6). The somewhat poor performance of the visual scores is related to the fact that almost 50% of the sample (i.e., 43% of males and 57% of females) was scored as “3”, evidencing a permanent difficulty in using an ordinal scale to visually classify a mastoid process as male or female. Yet, combining mastoid volume data with mastoid length or visual scores generally improves sex estimation compared to using volume data alone (Table 6), showing the usefulness of a multi-factorial approach also when there are clear correlations (around 0.5 – 0.75) between the different factors (Table 7).

These results show that mastoid volume measures can be valuable in a multi-factorial approach to sex estimation (Garvin and Klales, 2017; Langley et al., 2017), and even used alone if only fragmented parts of the temporal bone are available. The sex estimation accuracy of 70.6 % obtained for the Croatian population using Vol-2 is higher than the classifications achieved using traditional mastoid measurements for some African (Jaja et al., 2013), Brazilian (Galdames et al., 2008), Indian (Sumati et al., 2010), Saudi (Madadin et al., 2015), and white European and American (Bernard, 2008; Kemkes and Gobel, 2006; Spradley and Jantz, 2011) populations, but worse than the classification results achieved for Brazilian (de Paiva and Segre, 2003), other Indian (Gupta et al., 2012), Japanese (Nagaoka et al., 2008), Jordanian (Amin et al., 2015), and Thai (Manoonpol and Plakornkul, 2012; Sujarittham et al., 2011) groups. Because our traditional measurements showed that the SDI for the Croatian sample is lower than for most other groups, we speculate that sex estimation based on mastoid volumes may produce even better results for other populations.

Our comparison of the mastoid volumes 1, 2, and 3 illustrate the difficulties involved in defining the volume of a bony projection that lacks clear borders and nearby landmarks suitable for delineating it. Although Vol-2 appears to be the most promising volume for future applications, this volume could be improved by using alternative landmarks or/and planes for digital slicing of the mastoid, taking in consideration the developmental and functional features of the mastoid process, or/and relying on anatomical locations that can be easily and reproducibly identified in 3D models and that do not need to be strictly linked with the traditional landmark legacy (Wärmländer et al 2019). Interpopulation variability should be expected, as already indicated in Table 1 and 2. Volume analysis may of course perform differently in different populations. This may be most evident in the performance of volume 1, which uses landmarks and features known to differ between populations, such as opisthion, the cranial base chord, and foramen magnum (Zdilla et al., 2017).

Future research will likely be able to devise better metrics for mastoid volume measurements, which ideally should be tested on samples of known-sex individuals from different geographic regions and time periods (Petaros et al., 2017; Wärmländer and Sholts, 2011).

## Compliance with ethical standards

The 3D CT models were recorded in compliance with the current laws and ethical guidelines of the country where they were recorded.

